# The N-terminal GTPase of Miro1 regulates oligomer formation

**DOI:** 10.1101/2022.10.25.513744

**Authors:** Emily L. Eberhardt, Abdallah A. Mohamed, Seychelle Vos, Michael A. Cianfrocco

## Abstract

The outer-mitochondrial membrane protein Miro1 is critical for the regulation of mitochondrial trafficking. Miro1 contains two GTPase domains, where changes in the N-terminal GTPase nucleotide state strongly affects mitochondrial trafficking. Previous work showed that the GTP-locked mutation Miro1^P13V^ decreases trafficking and affects mitochondrial dynamics. Despite showing a clear role in Miro1 function, the molecular basis for this activity remains unknown. Using *in vitro* reconstitution, we demonstrate that Miro1 self-associates to form dimers and higher-ordered species. Structural characterization of Miro1^P13V^ suggests that the oligomers adopt a range of conformations *in vitro*. Additionally, Miro1^P13V^ has diminished interaction with its downstream cargo adapter TRAK1. These results indicate that the NGTPase of Miro1 facilitates the formation of higher-ordered species and suggests that changes in the oligomeric state may regulate mitochondrial trafficking through reduced association with TRAK1.

## INTRODUCTION

Microtubule-based trafficking of mitochondria is critical to fulfill the vast energy needs of cells. In heavily-branched, polarized cells such as neurons, this process must be tightly-regulated to ensure that mitochondria are present throughout the cell (Misgeld & Schwarz, 2017). Importantly, mitochondrial ATP production supports neuronal excitation, synaptic transmission, cell survival, and other cellular processes and must be regulated to ensure the proper distribution of mitochondria (Schwarz, 2013). At the core of mitochondrial trafficking is the motor-adaptor protein complex, consisting of the motor proteins kinesin-1 and cytoplasmic dynein-1 and cargo adaptor proteins that interact with cargo to promote bidirectional movement along microtubule filaments (Cason & Holzbaur, 2022). This process must be heavily-regulated, as dysregulation results in many disease pathologies such as Huntington’s disease, schizophrenia, amyotrophic lateral sclerosis, and Parkinson’s disease (Li *et al*, 2010; Norkett *et al*, 2016; Chen & Chan, 2009; Sheng & Cai, 2012; Rui *et al*, 2006; De Vos *et al*, 2007; Moon & Paek, 2015).

At the core of mitochondrial transport is the small G protein, *mi*tochondrial *r*h*o* (Miro), which interacts with the cargo adaptor TRAK to activate motor proteins (MacAskill *et al*, 2009; Guo *et al*, 2005; Fransson *et al*, 2006). Over the past twenty years, Miro1 has emerged as a critical player in mitochondrial trafficking. Miro1 is a surface-exposed, outer-mitochondrial membrane protein anchored to the mitochondria through a small C-terminal transmembrane tail (Fransson *et al*, 2003, 2006; Eberhardt *et al*, 2020). Interestingly, Miro1 contains two distinct, unique GTPase domains, which flank two calcium-binding EF hands. As a small GTPase, Miro1 cycles between a GTP-bound, active form and an inactive, GDP-bound form (Peters *et al*, 2018; Koshiba *et al*, 2011). Meanwhile, the EF-hands of Miro1 act as calcium sensors and arrest mitochondrial trafficking in the presence of increased local cellular calcium (Wang & Schwarz, 2009; Macaskill *et al*, 2009; Saotome *et al*, 2008).

Mutations that alter the GTP state of Miro1 have pronounced effects on mitochondrial trafficking and morphology. The N-terminal GTPase (NGTPase) is critical for the regulation of mitochondrial trafficking, although the effect of either the GTP- or the GDP-bound states is conflicting (Miro1^P13V^ and Miro1^T18N^, respectively) (Babic *et al*, 2015; Fransson *et al*, 2006; MacAskill *et al*, 2009). Some studies have suggested that Miro1^P13V^ inhibits mitochondrial trafficking (MacAskill *et al*, 2009), while others have suggested that it has less of an effect than the Miro1^T18N^ mutation (Babic *et al*, 2015; Davis *et al*, 2022; Norkett *et al*, 2020). Regardless, the GTP-bound state of Miro1 alters both mitochondrial dynamics and distribution and affects TRAK1 recruitment to mitochondria, suggesting that changes to the NGTPase affect proper mitochondrial activity (MacAskill *et al*, 2009; Babic *et al*, 2015; Fransson *et al*, 2003, 2006; Davis *et al*, 2022).

One potential mechanism by which Miro1 regulates mitochondrial trafficking is through Miro1 self-association. Miro1 forms discrete clusters through interaction with itself on the outer-mitochondrial membrane (Modi *et al*, 2019). Additionally, overexpression and subsequent immunoprecipitation of Miro1 demonstrated that Miro1 can co-immunoprecipitate itself in cell lysates (Kanfer *et al*, 2015; Nemani *et al*, 2018; Modi *et al*, 2019). These studies indicate a potential regulatory mechanism by which multiple Miro1 interact to form a complex that can act as a scaffold for motors and related machinery, such as DISC1 and Mitofusins (Norkett *et al*, 2020, 2016; Misko *et al*, 2010; Modi *et al*, 2019).

To recruit motor proteins to mitochondria, Miro1 associates with a member of the TRAK family of cargo adaptors (Fenton *et al*, 2021; Mitchell *et al*, 2021; Canty *et al*, 2021; Glater *et al*, 2006; van Spronsen *et al*, 2013; Randall *et al*, 2013; Henrichs *et al*, 2020; Brickley & Stephenson, 2011; Brickley *et al*, 2005; Stowers *et al*, 2002). Previous studies have demonstrated that Miro1 interacts with either one of the TRAK homologs, TRAK1 or TRAK2. In particular, the N-terminus of TRAK mediates interaction with motor proteins (van Spronsen *et al*, 2013), while the Miro-binding domain (TRAK2^476-700^/TRAK1^489-699^) has been proposed to mediate interaction with Miro1 (MacAskill *et al*, 2009).

Here, we investigated the effects of mutating the NGTPase of Miro1 to a GTP-like state (Miro1^P13V^). Using purified proteins, we demonstrate that Miro1^P13V^ promotes Miro1 self-association. Next, using negative stain electron microscopy, we show that Miro1 self-associates in multiple conformations, indicating that multiple interaction points within Miro1 govern self-association. Finally, we demonstrate that TRAK1 contains two binding sites for Miro1, one within TRAK1^395-532^ and another within TRAK1^562-635^. Miro1^P13V^ binds our TRAK constructs with reduced affinity, suggesting that the NGTPase affects trafficking through altered binding to TRAK1.

## RESULTS

Previous studies have indicated that the GTP-state of the NGTPase of Miro1 is critical for proper mitochondrial trafficking (Babic *et al*, 2015; Davis *et al*, 2022; MacAskill *et al*, 2009; Fransson *et al*, 2003, 2006). To determine the effect of stabilizing the NGTPase in a GTP-locked conformation, we introduced the point mutation P13V to the NGTPase (**Figure 1A**). After purification of recombinant Miro1^P13V^, we measured the effect of the mutation via size-exclusion chromatography (SEC) profile versus Miro1^WT^. As expected, Miro1^WT^ elutes as a single peak on SEC and runs as a single band on SDS-PAGE (**Figure 1B & Supplemental Figure 1A**). Surprisingly, the introduction of the P13V mutation affects the SEC profile, where Miro1^P13V^ runs as two distinct peaks. SDS-PAGE analysis shows a single band, indicating that both peaks of the SEC profile correspond to Miro1^P13V^ (**Figure 1C & Supplemental Figure 1B**). We can conclude that Miro1^WT^ and Miro1^P13V^ are not aggregating due to the lack of protein eluting in the void volume and the presence of discrete peaks at a late retention volume. These results suggest that Miro1^P13V^ induces the formation of higher-ordered species of Miro1, mediated by the NGTPase state.

**Figure 1.**
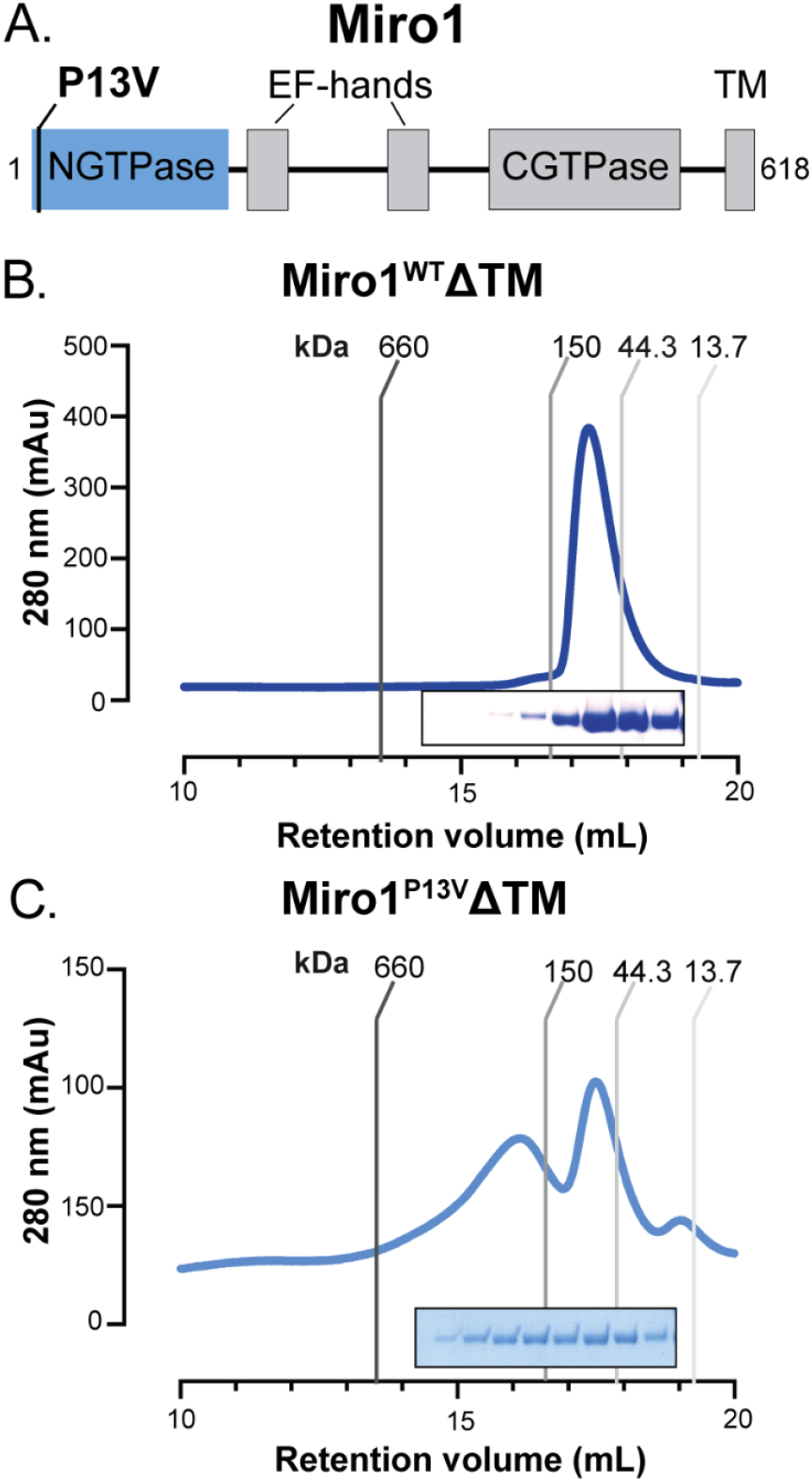
Mirol^P13V^ promotes the formation of higher-ordered Mirol complexes. (A) Domain architecture of Mirol protein. “P13V” denotes location of point mutation. Size-exclusion chromatography elution profile for purified (B) Miro1^WT^ΔTM and (C) Miro1^p13V^ ΔTM. For both (B) & (C) SDS-PAGE insets correspond to peak fractions (left to right). Void volume elutes before 10 mL.

### Miro1^P13V^ forms dimers and oligomers

To determine the molecular weight of the higher-ordered Miro1 species, we used size-exclusion chromatography multi-angle light scattering (SEC-MALS) to characterize Miro1^WT^ and Miro1^P13V^. Consistent with the results of SEC (**Figures 1B & 1C**) and the expected molecular weight of 67.86 kDa, Miro1^WT^ elutes in a single peak with a measured molecular weight of 75.51 kDa (**Figure 2A**). Meanwhile, Miro1^P13V^ migrates across multiple fractions in both monomeric (65.49 kDa) and dimeric peaks (137.46 kDa) (**Figure 2B**). In addition to the dimeric pool, larger complexes are eluting in the shoulder of the first peak. Due to poor separation of peaks in the shoulder, molecular weights could not be reliably measured in this region.

**Figure 2.**
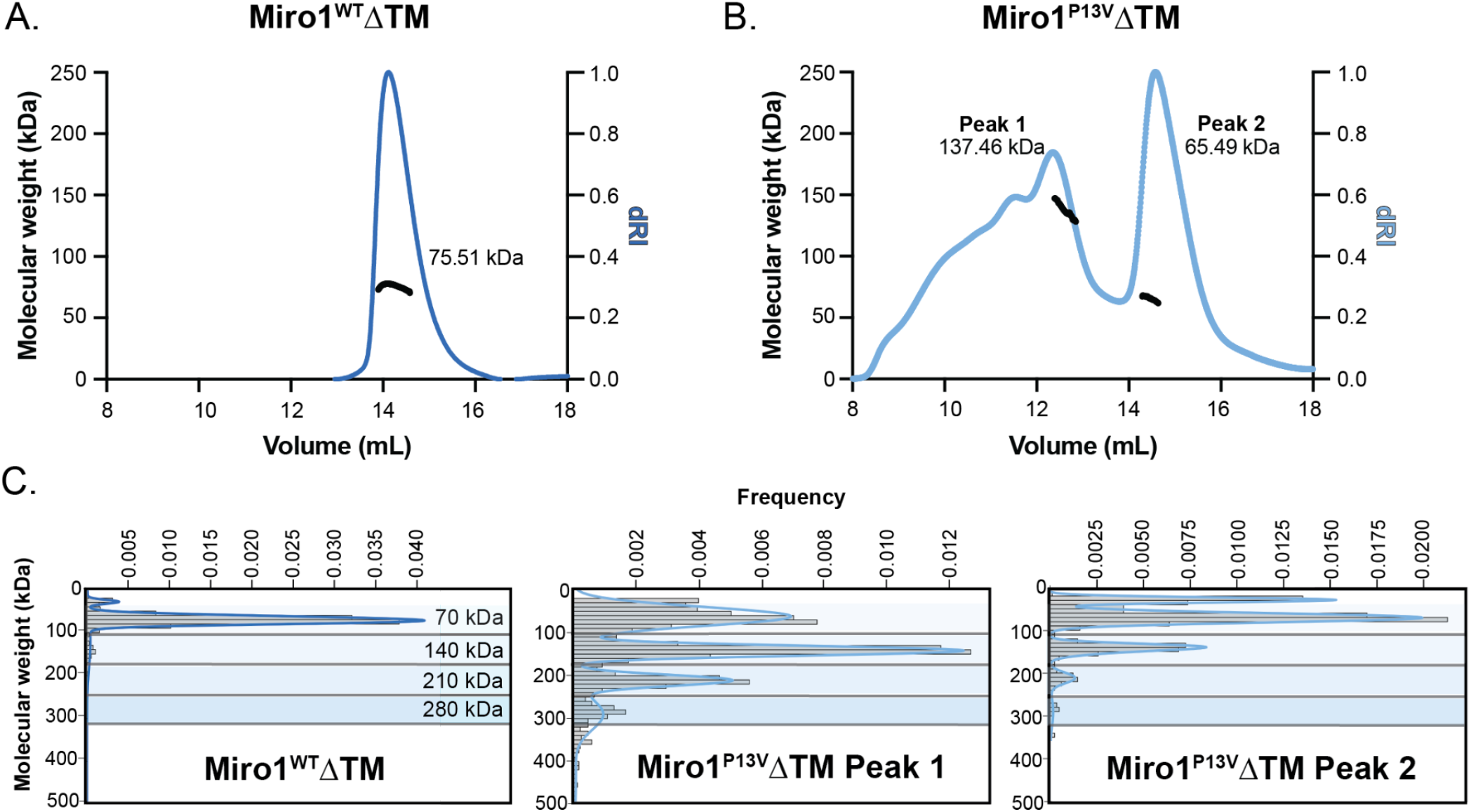
Miro1^P13V^ ΔTM forms dimers and oligomers. (A) SEC-MALS elution profile for Miro1^WT^ΔTM. (B) SEC-MALS elution profile for Miro1^P13V^ΔTM. (C) Mass photometry data showing mass distribution for Miro1^WT^ΔTM, Miro1^P13V^ΔTM Peak 1, and Miro1^P13V^ΔTM Peak 2.

To characterize the molecular weight of the higher-ordered species of Miro1^P13V^, we performed mass photometry. This allowed for higher-resolution characterization of species hidden in the long shoulder of the elution profile for Miro1^P13V^ (**Figure 2B**). Fractions for mass photometry were selected from the center of each SEC peak from both Miro1^WT^ and Miro1^P13V^. As expected, analysis of Miro1^WT^ showed a single species at ~72 kDa (**Figure 2C, left panel**). Surprisingly, the peak 1 fraction for Miro1^P13V^ showed Miro1 dimers, trimers (212 kDa), and tetramers (283 kDa) (**Figure 2C, middle panel**). Meanwhile, the peak 2 fraction for Miro1^P13V^ primarily formed monomers (72 kDa), with a minority (3%) forming dimers (138 kDa) (**Figure 2C, right panel**). These data indicate that the NGTPase mediates the formation of Miro1 oligomers.

### Miro1 NGTPase mediates an array of structural conformations

Next, we sought to characterize the impact of the P13V mutation on the structure of Miro1. We performed negative stain electron microscopy for peak fractions from Miro1^WT^ and Miro1^P13V^. At the micrograph level, Miro1^WT^ showed monomeric, monodisperse particles, whereas Miro1^P13V^ peak 1 was visibly clustered (**Supplemental Figures 2A & 2B**). To obtain more detailed structural information, we performed single particle analysis on collected datasets of each sample. For Miro1^WT^, monomeric Miro1 adopts a single conformation that appears as a ‘hockey stick’ with a bent domain extending off of a straight rod (**Figure 3A**). Meanwhile, analysis of the Miro1^P13V^ Peak 2 (low molecular weight) fraction showed a similar overall view of Miro1 compared to Miro1^WT^, with a small fraction appearing as dimers (**Figure 3B, top panel**). In the dimeric class average, Miro1^P13V^ adopts a partially folded-up conformation rolled up into a ball-like structure, with part of the domain architecture hanging off.

**Figure 3.**
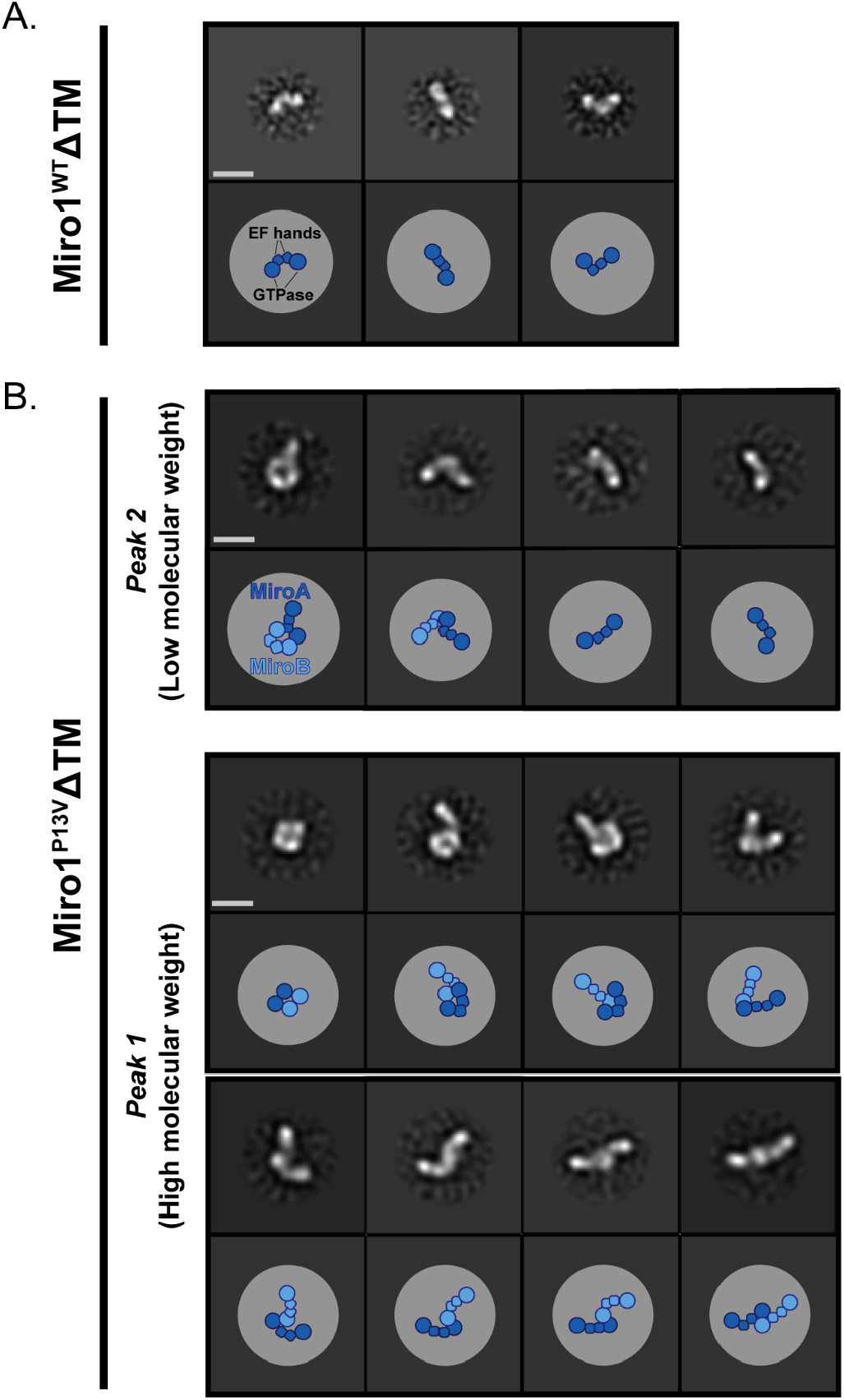
The Mirol NGTPase mediates multiple dimer conformations. 2D classes and corresponding cartoon diagrams depicting potential organizations for (A) Miro1^WT^ and (B) Miro1^P13V^. Scale bar represents 100 Å.

In contrast to Miro1^WT^ and Miro1^P13V^ Peak 2, Peak 1 (high molecular weight) Miro1^P13V^ showed that Miro1 forms an array of dimeric conformations. In addition to the rolled-up conformation of Miro1 and potential top-down views (**Figure 3B, middle panel**), Miro1^P13V^ appears to form multiple GTPase-GTPase interactions. The multiple conformations suggest that there are a vast array of interactions that occur between GTPases. Additionally, there may be conformations of Miro1^P13V^ where both the GTPase and an EF-hand interact (**Figure 3B, middle and bottom panels**). These 2D class averages suggest that the NGTPase of Miro1 is responsible for forming an array of Miro1 self-associations.

### Miro1 NGTPase state affects binding to TRAK1

Interaction between Miro1 and TRAK1 is at the center of the motor-adaptor complex and is necessary for mitochondrial trafficking (Guo *et al*, 2005; Glater *et al*, 2006; van Spronsen *et al*, 2013; Fransson *et al*, 2006; Canty *et al*, 2021; Henrichs *et al*, 2020; Fenton *et al*, 2021). Since mutation of the NGTPase results in defects in mitochondrial trafficking and function, we decided to test if this phenotype is due to decreased affinity of TRAK1 for Miro1. Therefore, we performed a set of co-immunoprecipitation experiments with over-expressed recombinant Miro1 and TRAK1 fragments. We then measured binding between Miro1 and TRAK1 by determining protein binding to beads using SDS-PAGE. Based on the previously described Miro-binding domain, we generated a set of truncations within the TRAK1 amino acid residues 489-699 to determine the minimal region on TRAK1 for interaction with Miro1 (**Figure 4A**). Of the minimal fragments generated, TRAK1^562-635^ was robustly overexpressed and used for subsequent studies. Interestingly, Miro1 was able to pull-down overexpressed TRAK1^562-635^. However, there is a slightly lower affinity for TRAK^562-635^ with Miro1^P13V^ compared to Miro1^WT^ (**Figure 4C**). Quantification of the band intensity of Miro1^P13V^ suggests a ~30% reduction in TRAK1^562-635^ interaction (*p* = 0.0136) (**Figure 4F**).

**Figure 4.**
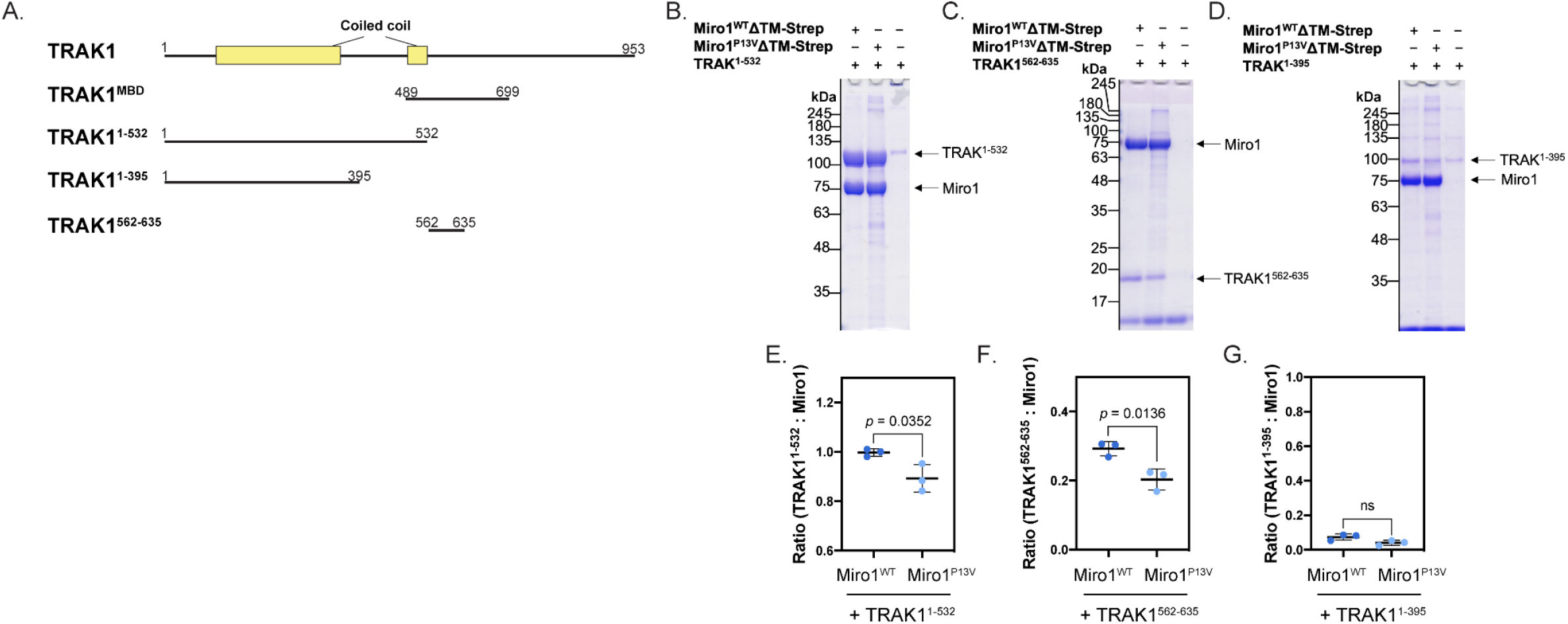
Miro1^P13V^ has a reduced affinity forTRAKI. (A) TRAK1 domain architecture, TRAK1^Mira-binding domain^, and TRAK1 constructs used. Strep-bead pull-down in the presence of Miro1^WT^-Strep, Miro1^P13V^-Strep, or buffer control and (B) TRAK1^1-532^, (C) TRAK1^562-635^, or (D) TRAK1^1-395^. Quantification of the band intensity for the ratio of (E) TRAK1^1-532^ : Mirol-Strep (*p* = 0.0352 (Student’s t-test)), (F) TRAK1^562-635^ : Mirol-Strep (*p* = 0.0136 (Student’s t-test)), and (G) TRAK1^1-395^ : Mirol-Strep (*p >* 0.05 (Student’s t-test)).

To determine if Miro1 interacts with any other regions of TRAK1, we utilized a TRAK1^1-532^ construct. Surprisingly, Miro1^WT^ and Miro1^P13V^ also interacted with TRAK1^1-532^ (**Figure 4B**). Analysis of the interaction also showed a slight decrease in affinity for Miro1^P13V^ with TRAK1^1-532^ as compared to Miro1^WT^ (*p* = 0.0352) (**Figure 4E**).

To rule out potential non-specific interaction between Miro1 and the TRAK1 constructs, we next tested the interaction between Miro1 and TRAK1^1-395^, which contains the region where motor proteins bind (Randall *et al*, 2013; Canty *et al*, 2021; Henrichs *et al*, 2020; Fenton *et al*, 2021). TRAK1^1-395^ interacted with Miro1^WT^, Miro1^P13V.^, and the negative control with a similar affinity, suggesting minimal to no interaction with TRAK1^1-395^ (**Figures 4D & 4G**). Taken together, these results suggest that Miro1 binds to TRAK1 using two binding sites that are each sensitive to the NGTPase GTP state.

## DISCUSSION

Here, we describe the molecular mechanism by which the GTP-locked NGTPase affects Miro1 self-association. We utilized recombinant Miro1^WT^ and Miro1^P13V^ to demonstrate that when the NGTPase is locked in the GTP-bound state, Miro1 self-associates into discrete oligomeric species. Further, we show Miro1^WT^ binds to TRAK1 using two distinct regions on TRAK1, and Miro1^P13V^ has reduced binding to these sites. Combined, these findings suggest that the Miro1 NGTPase mediates self-association, which diminishes association with the cargo adaptor, TRAK1.

From yeast to *Drosophila* to mammalian cells, mutations in the Miro1 GTPase domains demonstrate a critical role for the NGTPase in mitochondrial trafficking and activity (Frederick *et al*, 2004; Fransson *et al*, 2003; MacAskill *et al*, 2009; Koshiba *et al*, 2011; Babic *et al*, 2015; Wang & Schwarz, 2009). Meanwhile, mutation of the CGTPase appears to have little to no effect on mitochondrial dynamics or trafficking (Babic *et al*, 2015; Fransson *et al*, 2003, 2006; MacAskill *et al*, 2009; Davis *et al*, 2022). In particular, the Miro1^P13V^ mutation results in mitochondrial aggregation, trafficking defects, and cellular apoptosis, although the mechanism by which this occurs has remained unknown (Fransson *et al*, 2003, 2006; Babic *et al*, 2015; MacAskill *et al*, 2009; Davis *et al*, 2022; Norkett *et al*, 2020). While these studies have provided broad insight into the widespread cellular defects incurred by Miro1^P13V^, we suggest that these phenotypes could be explained by Miro1 self-association, where oligomeric Miro1 may change binding partners or cause membrane-membrane contacts. We propose that Miro1 self-association is a vital regulatory process for mitochondrial trafficking as well as overall mitochondrial function.

Using purified, recombinant proteins, our data show that Miro1 is sufficient for self-association. This finding is supported by previous studies, where Miro1 clusters (Nemani *et al*, 2018; Modi *et al*, 2019) and immunoprecipitates itself (Kanfer *et al*, 2015; Modi *et al*, 2019). However, since these studies were performed using cell lysates, other proteins may be mediating the interaction between Miro1 proteins. Our data support previous findings and suggest that these self-associations are functional, due to the lack of aggregation seen in our biochemical analysis. Finally, since our data demonstrate that purified, recombinant Miro1 self-associates, we can conclude that no other proteins are required to facilitate the interaction.

Previous structural data suggest that GTPase domains of Miro1 (NGTPase or CGTPase alone) dimerize along a non-crystallographic plane (Klosowiak *et al*, 2013; Smith *et al*, 2020; Klosowiak *et al*, 2016). However, due to the tight coupling of the CGTPase to the EF-hands, the CGTPase is incapable of mediating dimerization (Klosowiak *et al*, 2016). Meanwhile, the NGTPase does not form a stable interface with the rest of the protein, suggesting conformational freedom to interact with itself or other proteins (Smith *et al*, 2020). This data is supported by accessible hydrophobic patches on the NGTPase that suggest that Miro1 can coordinate associations with other proteins (Smith *et al*, 2020). Indeed, Miro1 interacts with multiple dimeric, downstream binding partners, including Mitofusins, Drp1, and DISC1 (Norkett *et al*, 2020; Daumke & Roux, 2017; Macdonald *et al*, 2014). Additionally, some of these binding partners specifically interact with Miro1^P13V^ (Norkett *et al*, 2020; Kanfer *et al*, 2015). Therefore, these findings suggest that Miro1 oligomers coordinate interaction with binding partners through the NGTPase, where Miro1 can interact with itself or other proteins when GTP-bound.

We propose that Miro1 is able to interact with itself laterally across the outer-mitochondrial membrane and with Miro1 proteins on other organelles. Our model helps to understand how Miro1 forms clusters on the outer-mitochondrial membrane (Modi *et al*, 2019) and also exists at multiple organelle contact points, including the ER-mitochondrial contact sites (Modi *et al*, 2019; Lee *et al*, 2016) and at peroxisomes (Covill-Cooke *et al*, 2020; Castro *et al*, 2018). The multiple 2D conformations we visualize for Miro1 interaction could explain how Miro1 self-associates depending on its spatiotemporal location.

Our data suggest that Miro1 self-association may diminish interaction with TRAK1, due to steric or allosteric hindrance. Compared to Miro1^WT^, Miro1^P13V^ has decreased interaction with two TRAK1 binding sites, mediated by the GTP-state of the NGTPase. While previous data have suggested that Miro1 interacts with TRAK^489-699^, we discovered that this region contains two independent binding sites, although there may be more interaction sites within this region (MacAskill *et al*, 2009; Mitchell *et al*, 2021). Since both sites appear sensitive to the NGTPase state, this could suggest that mitochondrial trafficking is partially mediated by GTPase cycling of Miro1. Further investigation into minimal binding regions will provide much-needed insight into how Miro1 binds TRAK1 and how this association changes with NGTPase state.

## ACKNOWLEDGEMENTS

We are exceptionally thankful for the feedback on this manuscript by Dr. Thomas L. Schwarz, Dr. Morgan E. DeSantis, Tyler M. Hoard, and Hye Jee (Lily) Hahn. We would like to thank Jaykrishnan Nandakumar, Cassandra Zuckerman, and Ritvijia Agrawal for their assistance in running SEC-MALS experiments and data analysis. We would also like to thank Zhenyu Tan for supplying plasmids. The research reported in this publication was supported by the University of Michigan Cryo-EM Facility (U-M Cryo-EM). U-M Cryo-EM is grateful for support from the U-M Life Sciences Institute and the U-M Biosciences Initiative. This work was supported by the National Institutes of Health grant S10OD020011. A.M. is supported by funding from the NSF-GRFP Fellowship #1745302.

## Author Contributions

E.L.E performed all biochemical and structural experiments. E.L.E and M.A.C performed negative stain analysis. A.M. and S.V. performed mass photometry experiments. Z.T. generated Miro^WT^, TRAK^1-395^, and TRAK^1-532^ constructs. E.L.E and M.A.C prepared this manuscript with comments from all authors.

## METHODS

### Constructs

The constructs used in this study were:

| Name | Plasmid Details |
| --- | --- |
| Miro1<sup>WT</sup> | pFastBac-His-ZZ-TEV-Miro1<sup>1-592</sup> and pACEBac-Miro1<sup>1-592</sup>-TEV-Strep |
| Miro1<sup>P13V</sup> | pFastBac-His-ZZ-TEV-Miro1<sup>1-592ΔP13V</sup> and pACEBac-Miro1<sup>1-592ΔP13V</sup>-TEV-Strep |
| TRAK1<sup>1-532</sup> | pACEBac-His-ZZ-TEV-SNAP-TRAK1<sup>1-532</sup> |
| TRAK1<sup>562-635</sup> | prSET-His-GB1-TEV-TRAK1<sup>562-635</sup> |
| TRAK1<sup>1-395</sup> | pACEBac-His-ZZ-SNAP-TRAK1<sup>1-395</sup> |

### PCR Cloning

ZZ-tag-TEVpRSET-A-Miro1^P13V^ was purchased from Thermo Scientific™ using the GeneArt tool. The Miro1^P13V^ gene was then inserted into pFastbac for insect cell expression using HiFi DNA Assembly.

The pFastbac vector was PCR amplified using forward and reverse primers: 5’-AGCACCTTTTAATAAGAATTCAAAGGCCTACGTCGAC-3’ and 5’-CTTTTTTCATTGAGCCCTGGAAGTACAGGT-3’

The Miro1^P13V^ gene was PCR amplified using: 5’-CAGGGCTCAATGAAAAAAGATGTGCGTATTCTGCTGGT-3’ and 5’-TTTGAATTCTTATTAAAAGGTGCTGCTTTTCAGATCTG-3’

Strep-tagged Miro1 constructs were generated from a PCR amplification of Strep-tag containing backbone: 5’-GCACCTTTGAGAACCTGTACTTCCAGTCTGGTTCTGG-3’ and 5’-TTTTTCATGGTGGCGGATCCGCG-3’ and Miro1 genes: 5’-CCGCCACCATGAAAAAAGATGTGCGTATTCTGCTGG-3’ and 5’-CAGGTTCTCAAAGGTGCTGCTTTTCAGATCTG-3’

GB1-His backbone was generated using 5’-CAGGTGATCATAATATTGGAAGTGGATAACGGATCCGAA-3’ and 5’-GATGAAAAAGAGGATTGGAAGTACAGGTTCTCGG-3’

TRAK^562-635^ fragment was generated using 5’-CTTCCAATCCTCTTTTTCATCACGTTCTTATCTGCCAGAAA-3’ and 5’-ACTTCCAATATTATGATCACCTGTATCGTCCTCTTCAAAAT-3’

### Insect Cell Protein Purification

The baculovirus expression system was used to express Miro1^WT^, Miro1^P13V^, TRAK^1-395^, and TRAK^1-532^. Briefly, the Bac-to-Bac system (Life Technologies) was used to generate recombinant baculovirus in insect cells. High Five™ cells (Gibco™), grown to 1.6-2 million cells/ml in Insect-XPRESS media (Lonza™), were infected with P1 viral stocks at 16 ml/L. *Spogopteria fruteria* (Sf9) or High Five™ cell pellets were collected 48-72 hours post-infection. Frozen cell pellets were resuspended in Miro lysis buffer [30 mM HEPES (pH 7.4), 150 mM KAc, 2 mM MgAc, 3 mM CaCl_2_, 0.1 mM GTP, 5% glycerol, 1 x protease inhibitor cocktail–Sigma] or TRAK lysis buffer (Miro lysis buffer without CaCl_2_ or GTP) lysed using a dounce homogenizer, fitted with a tight pestle, with 20 strokes. Two microliters of benzonase nuclease (MilliporeSigma) were added to the samples before they were ultracentrifuged at 40,000 x g for 35 minutes at 4°C. At this point, if the clarified lysate was to be used in downstream pull-down assays, the clarified lysate was flash frozen using LN_2_. Otherwise, the resulting clarified lysate was batch-bound to ~1 mL of settled IgG sepharose (GE Healthcare) equilibrated in lysis buffer and allowed to incubate on a roller for 2 hours at 4°C. Protein was eluted from sepharose by cleavage with Tobacco Etch Virus (TEV) protease at 16°C for 2 hours. The eluate was separated from the resin using an Ultrafree-CL centrifugal filter (MilliporeSigma) and spun at 1,000 x g for 2 minutes. The eluate was then concentrated to ~1-2 mg/mL in a 10k MWCO Amicon spin filter (MilliporeSigma). TEV and residual affinity tag was removed by passing the protein through a Superdex 200 Increase 10/300 GL (GE Healthcare) in SEC buffer (30 mM HEPES pH 7.4, 150 mM KAc, 2 mM MgAc, 3 mM CaCl2, 0.1 mM GTP). Fractions were assessed by mixing twelve microliters of fraction with 4 microliters of 4X NuPAGE™ LDS sample buffer (Invitrogen™), boiling, and loading 15 microliters onto SDS-PAGE. Protein concentration was assessed by staining the gel with Imperial™ Protein Stain (Thermo Scientific™). Fractions were pooled at flash frozen in liquid nitrogen before storage at -80°C for further use.

### SEC-MALS

For Miro1^WT^, protein samples (stored at -80°C) purified using the protocol in “**Insect Cell Protein Purification**” were thawed on ice and pooled to 500 μL of ~1 mg/mL. For Miro1^P13V^, fresh protein was concentrated to 500 μL at ~1 mg/mL. Samples were injected onto a Superdex Increase 200 column (GE Healthcare) in line with a DAWN HELIOS II MALS detector (Wyatt Technology) and an Optilab T-rEX differential refractometer (Wyatt Technology). Differential refractive index and light scattering data were measured and analyzed using ASTRA 6 software (Wyatt Technology). Molecular weights were calculated via extrapolation from Zimm plots using a d*n*/d*c* value of 0.185 ml/g. Finally, molecular weight outputs were aligned with retention volume in Microsoft Excel before data was plotted in Graphpad Prism.

### Mass photometry

Using the Refeyn Two^MP^ (Refeyn), coverslips were prepared by washing one time with filtered H_2_O and then twice with filtered 20% ethanol. In between washes, the coverslip was dried with compressed air. Gaskets (MillliporeSigma) were then placed on the cleaned coverslip. Next, a drop of optical oil was added to the Refeyn 2 optic, before placing the coverslip with gasket on the stage and centering the laser with the gasket opening. Then, 15 μL of 1 x PBS was added to a gasket and autofocused using the AcquireMP software. Next, NativeMark™ Standard (Invitrogen™) was diluted 1:50, before adding 5 μL to the PBS. The coverslip was then moved to the next open gasket. Miro1 samples were diluted to 10 nM in 1 x PBS, and 20 μL was added to open gasket holes. Movies were acquired for each sample using AcquireMP software. Data was exported as a .csv file and analyzed by applying Gaussian distributions to each peak.

### Negative Stain Electron Microscopy

For negative stain electron microscopy, Miro1 samples were diluted to ~0.01-0.05 mg/mL in SEC buffer. Formvar, carbon-coated copper grids were glow discharged with a PELCO easiGlow for 30 s at 5 mA. Four microliters of each Miro1 sample were applied to the carbon face of the grid for one minute. The grid was then wicked using a small piece of Whatman blotting paper, then placed face down in ∼30 μL of distilled water and wicked, two times, before incubation in 0.075% uranyl formate solution three times for 1 second, 10 seconds, and finally 45 seconds. The grid was then wicked with filter paper and allowed to dry.

### Micrograph processing and 2D class averages

Grids were screened at 22,000x magnification using a Morgagni (FEI) transmission electron microscope and imaged with a CCD camera (2.1 Å/pixel @ 22,000x). Datasets were collected using a Tecnai-12 (FEI) at 42,000x magnification (1.45 Å pixel size). The resulting micrographs were collected with Leginon (Suloway *et al*, 2005) and processed using RELION (Kimanius *et al*, 2021). First, CTFFIND4.1 was used to estimate the CTF for each micrograph. Micrographs with CTF estimations higher than 14 Å were thrown out. Particles were picked using the Laplacian-of-Gaussian picker (180Å - 220Å) and adjusted default threshold (stddev) set to 1 with otherwise default parameters. The final numbers of micrographs and particles for each dataset were as follows: Miro1^WT^ - 66 micrographs and 11,789 particles, Miro1^P13V^ Peak 1-28 micrographs and 5485 particles, and Miro1^P13V^ Peak 2-24 micrographs and 5399 particles. Particles were extracted using a 300 pix box size and rescaled to 100 pixels. Next, 2D class averages were generated using 35 iterations, 250 Å mask diameter (200 Å for Miro1^WT^), and all other default parameters. “Good” class averages (visualized by expert) were selected for another round of 2D classification and subsequently characterized by visual inspection.

### TRAK^562-635^ Protein Purification

BL21 *E. coli* were transfected using the High-Efficiency Transformation Protocol (NEB). Single colonies were selected from carbenicillin LB plates and plated into 5 mL starter cultures, grown at 37°C, 220 rpm for 4-7 hours. Starter cultures were then added to 1 Liter of Hyperbroth media (Athena Enzyme Systems) with supplemented glucose nutrient and carbenicillin at 37°C, 220 rpm for 4-6 hours until >1 OD. The temperature was then lowered to 20°C before the addition of 1M IPTG and shaken for another 18-22 hours. Cell material was centrifuged at 4°C at 3,000 x g for 15 minutes, washed with 1X PBS, and spun for an additional 15 minutes at 4°C at 3,000 x g. Pellets were then scraped into a conical tube and flash-frozen using LN_2_. To generate cell lysates, cell pellets were thawed on ice before the addition of TRAK buffer [30 mM HEPES (pH 7.4), 150 mM KAc, and 2 mM MgAc, 0.1 mM GTP, 1 mM DTT, 5% glycerol]. Cell suspensions were sonicated using 80% duty cycle, output tip at 8, for 30-second intervals of sonication followed by 15-second stirring for a total of 8 minutes. The resulting lysate was ultracentrifuged at 40,000 x g for 35 minutes at 4°C. Clarified lysate was flash-frozen using LN_2_ and stored at 80°C until further use.

### *In vitro* pull-down assays

Per condition, one hundred microliters of settled Streptactin XT 4Flow resin (IBA) was equilibrated with TRAK buffer and 2 mLs of clarified Miro1-Strep lysate for 2 hours at 4°C. The Miro1-bound resin was then washed 4 times with 15 column volumes (CV) of TRAK buffer. A small sample of resin was added to 4x LDS buffer, boiled, and visually quantified by SDS-PAGE to determine the relative amount of Miro1-Strep TRAK to Streptactin resin. To control for nonspecific TRAK to the resin, 100 microliters of resin was incubated in the absence of Strep-tagged protein, before addition of two mLs of ZZ or His-tagged TRAK (TRAK^1-532^, TRAK^1-395^, TRAK^562-635^). Upon addition of secondary protein lysate, the mixture was incubated for 2 hours at 4°C. The resin was then washed 4 times with 15 CV of TRAK buffer. After the final wash, each sample was resuspended in 100 μL of 2x NuPAGE™ LDS sample buffer (Invitrogen™) in TRAK buffer and boiled. Ten microliters of the supernatant were loaded onto an SDS-PAGE gel. Gels were stained with Imperial™ Protein Stain (Thermo Scientific™), and band intensities were quantified with ImageJ. Each experiment was performed in triplicate.

### Densitometry Analysis

Gel images were opened in the ImageJ software suite, inverted, and converted to 16-bit. To equalize all signal, the contrast was set to a minimum of 0 and a maximum of 200 for all gels. Gels were straightened, and individual lanes were assigned for regions containing both bands. The area under each curve was measured by drawing a line under each intensity curve and measuring the area using the magic wand tool. Negative control values were subtracted from experimental conditions, and the ratio of TRAK1 to Miro1 (on the bead) was calculated and plotted.

