## Supplemental figures 1&2 for "The N-terminal GTPase of Miro1 regulates oligomer formation"

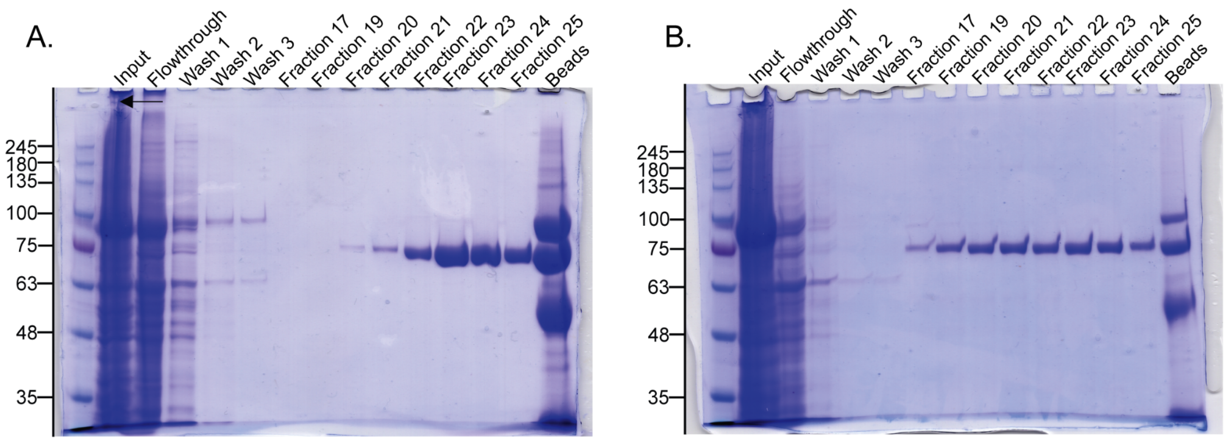


**Supplemental Figure 1. Original SDS-PAGE gel images.**
SDS-PAGE gels for (A) Miro1$^{WT}$ and (B) Miro1$^{P13V}$ preparation.

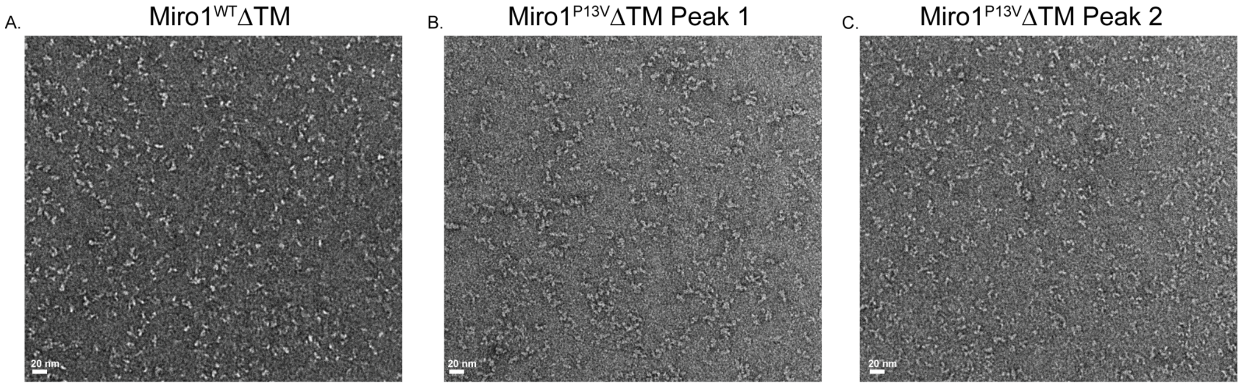


**Supplemental Figure 2. Representative negative stain micrographs of Miro1.**
Negative stain for (A) Miro1$^{WT}\Delta$TM, (B) Miro1$^{P13V}\Delta$TM single peak (Peak 1), and (C) Miro1$^{P13V}\Delta$TM single peak (Peak 2).
